# Power Calculator for Detecting Allelic Imbalance Using Hierarchical Bayesian Model

**DOI:** 10.1101/2021.07.10.451873

**Authors:** Katrina Sherbina, Luis G. León-Novelo, Sergey V. Nuzhdin, Lauren M. McIntyre, Fabio Marroni

## Abstract

Allelic imbalance (AI) is the differential expression of the two alleles in a diploid. AI can vary between tissues, treatments, and environments. Statistical methods for testing in this area exist, with impacts of explosive type I error in the presence of bias well understood. However, for study design, the more important and understudied problem is the type II error and power. As the biological questions for this type of study explode, and the costs of the technology plummet, what is more important: reads or replicates? How small of an interaction can be detected while keeping the type I error at bay? Here we present a simulation study that demonstrates that the proper model can control type I error below 5% for most scenarios. We find that a minimum of 2400, 480, and 240 allele specific reads divided equally among 12, 5, and 3 replicates is needed to detect a 10%, 20%, and 30%, respectively, deviation from allelic balance in a condition with power >80%. A minimum of 960 and 240 allele specific reads is needed to detect a 20% or 30% difference in AI between conditions with comparable power but these reads need to be divided amongst 8 replicates. Higher numbers of replicates increase power more than adding coverage without affecting type I error. We provide a Python package that enables simulation of AI scenarios and enables individuals to estimate type I error and power in detecting AI and differences in AI between conditions tailored to their own specific study needs.

## Introduction

Gene expression in a diploid individual is the result of the combined expression of both alleles. Allele Specific Expression (ASE) is the amount of mRNA transcribed at each allele. The two alleles of a diploid individual can show significantly different expression, a condition termed allelic imbalance (AI) [1]. AI is a result of genetic variation in regulation in *cis* (*e*.*g*. promoters, enhancers, and other noncoding sequences), *trans* (*e*.*g*. transcription factors) or resulting from *cis* by *trans* interactions [1–7]. AI has been observed as a consequence of imprinting [8–10] and nonsense mediated decay [11] and has been shown to contribute to heterosis [12] and hybrid incompatibility [13]. The extent of AI in human tissues can give information on the impact of heterozygous mutations on the expression of the mutated allele in healthy [14] or cancerous human tissues [15]. Also, loss of heterozygosity can be detected using AI [16, 17].

Comparing AI between conditions or tissues can provide new insights into the mechanisms of gene expression regulation [6, 9, 18–23]. Most often, these comparisons are heuristic without a formal statistical test. However, statistical comparisons have been made of heterogeneity in AI between mated and virgin *Drosophila* female head tissue [19], human tissues types within an individual [11, 22, 24], and cell subpopulations in different developmental stages [25]. Some statistical tests have been performed to assess whether *cis* effects differ among alleles in a population [7] or in parent of origin effects in mice [5].

Type I error in AI studies has been well explored and is known to be high, particularly when failing to account for map bias [26], and/or using the binomial test [5, 27–31]. What is currently absent from the literature is an understanding of the power for studies of AI and, in particular, what the best allocation of resources is for boosting power for detection of AI when the hypothesis of interest is a comparison of AI *between* conditions. What is more important: more reads or more replicates? Is there a minimum number of replicates needed? A minimum number of allele specific reads? As read numbers per sample are dropping in price with the capacity of the new technologies, and the per sample cost of library preparation dramatically lower than a decade ago, it is time to stop relying on the magic number 3 and determine the necessary size and scope of such studies to control type I *for* a particular type II error/power. It is common practice to assess power before embarking on association studies [32, 33], but no tool is currently available for assessing power for detecting AI and differences in AI between conditions.

To address this need, we present here the package BayesASE_power. It consists of tools to enable the user to simulate RNA-seq read counts under a previously published Bayesian model of AI [7, 28] with any number of replicates, reads, and AI. The results are aggregated across multiple simulated datasets to estimate Type I error and power. We demonstrate how to use BayesASE_power to plan experiments to achieve the desired power in detecting AI within a condition and/or interactions of AI between conditions.

## Methods

The model used for detection of AI in any condition and for comparing levels of AI between any two conditions has been described earlier and implemented in the package BayesASE [7, 19]. We give here the basic definitions and refer the reader to Supplementary Methods for further details.

One important parameter in determining AI is the probability *r* of a read aligning to allele *g1* (*g2*) given that it came from that allele, that we define as *r*_*i,g*1_ (*r*_*i,g*2_). Low values of these probabilities correspond to a high degree of ambiguously mapped reads, which occurs when there is little sequence divergence between the two alleles. Reads that do not map ambiguously are termed allele specific reads.

AI in condition *i* is measured by the parameter *θ*_*i*_ representing the proportion of reads originating from the allele *g1*, which that can be written as follows:

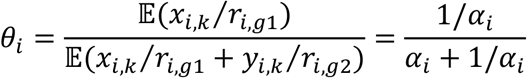

When *θ*_*i*_ is close to 0, we have one extreme case of AI with all the reads originating from *g2*. When *θ*_*i*_ *=* 0.5, we have perfect allelic balance with 50% of the reads from each allele. With *θ*_*i*_ *=* 1, we are in the opposite direction of extreme AI with all the reads originating from *g1*.

The following null hypotheses are defined:

1. Allelic balance in condition 1, *i*.*e*. null *H1*: *θ*_1_ *=* 0.5 or equivalently *α*_1_ *=* 1.
2. Allelic balance in condition 2, *i*.*e*. null *H2*: *θ*_2_ *=* 0.5 or equivalently *α*_2_ *=* 1.
3. Level of AI is the same in both conditions, *i*.*e*. null *H3*: *θ*_1_ *= θ*_2_ or equivalently *α*_1_ *= α*_2_.

To test these hypotheses, three cases are defined (Figure 1):

1. H1, H2 and H3 are satisfied
2. H1 is satisfied, H2 and H3 are violated
3. H1 and H2 are violated, H3 is satisfied

**Figure 1.**
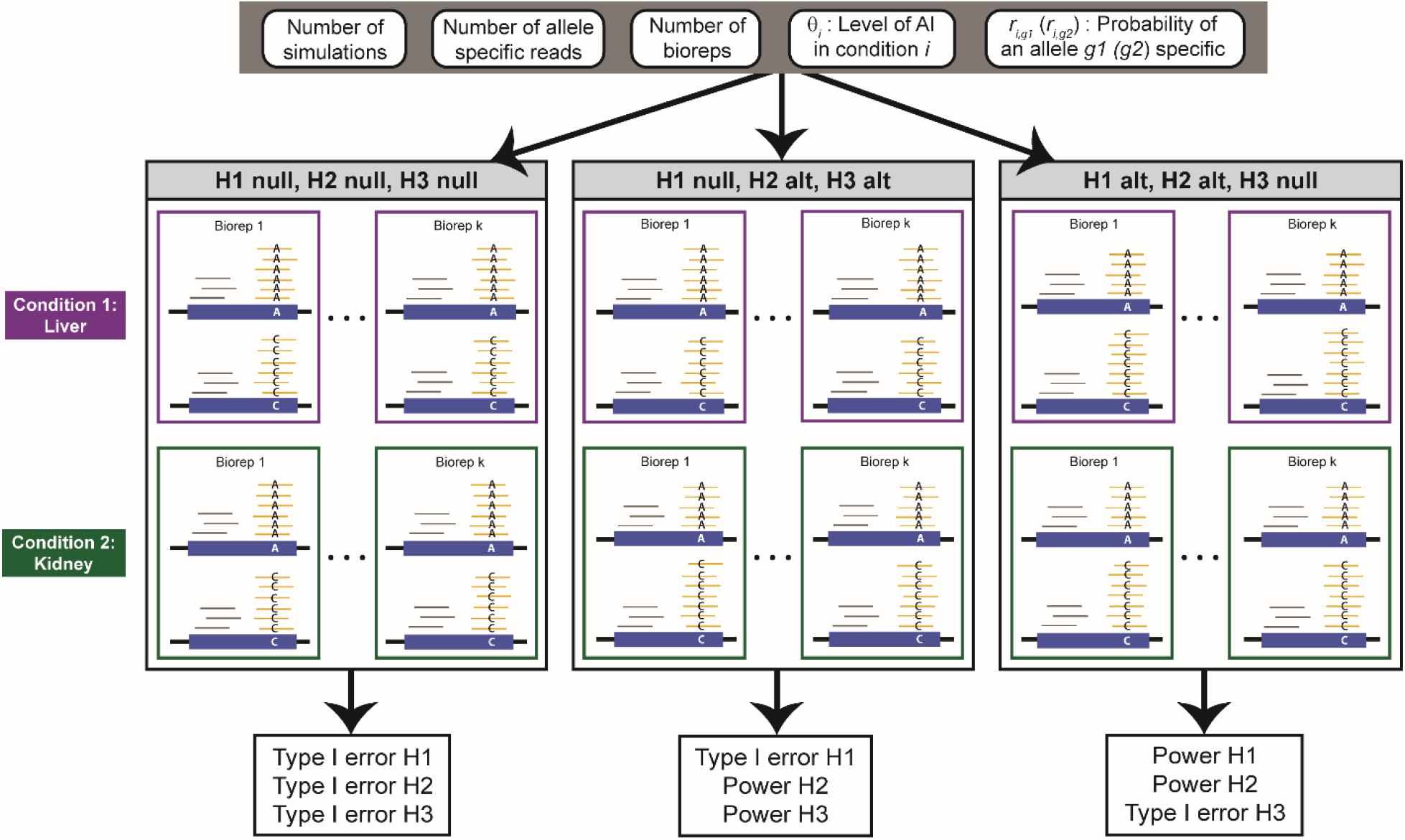
Read counts are simulated for different scenarios in two conditions. A scenario is defined as a specific number of simulations, number of allele specific reads, number of biological replicates (bioreps), level of allelic imbalance (AI) θ, and the probability of mapping an allele *g1* (*g2*) specific read. Without loss of generality, let allele *g1* be allele A and *g2* be allele C (blue boxes). The number of allele specific reads (yellow reads) is the sum of unambiguously mapped reads in the experiment. Grey reads are reads that map equally well, *i*.*e*. ambiguously, to both alleles. Biological replicates in an experiment are samples from the same genotype and condition. In this example, there are *k* biological replicates, 12 × *k* allele specific reads, and the probability of an allele specific read is *r*_*i,g*1_ *= r*_*i,g*2_ *=* 0.8. The null H1 and H2 hypotheses are allelic balance θ_1_=0.5 in condition 1 (ex. liver) and θ_2_=0.5 in condition 2 (ex. kidney), respectively. These cases are used to estimate the Type I error in rejecting allelic balance in conditions 1 (H1) and 2 (H2). In this example, θ_1_=0.55 under the alternative (alt) H1 hypothesis and θ_2_=0.55 under the alternative (alt) H2 hypothesis. These cases are used to estimate the power in rejecting allelic balance in conditions 1 (H1) and 2 (H2). θ_1_=0.5 and θ_2_=0.55 under the alternative (alt) H3 hypothesis, which allows estimation of the power rejecting equal levels of AI between the two conditions (H3). The null H3 hypothesis is simulated in both the complete null case: θ_1_= θ_2_=0.5 and in the scenario where there is allelic imbalance in both conditions θ_1_=θ_2_=0.55. These cases can be used to estimate the Type I error in rejecting equal levels of AI between the two conditions (H3).

In our simulation, magnitudes of deviation from the null are reported as Δ*AI*. Given *θ*_0_ *=* 0.5, 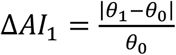 for H1, 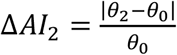 for H2, and 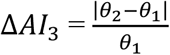 for H3. Simulated deviations of Δ*AI* from the null are moderate, generally between 0.1 and 0.3, with a maximum of 0.5.

Scenarios that vary the number of allele specific reads, number of replicates, and AI in the different cases were simulated (Table S1).

## Results

Type I error is controlled except when total allele specific reads exceed 2400 allele specific reads dispersed across 8 or more biological replicates (Fig 2a-2b). However, type I error is always less than 0.08.

**Figure 2.**
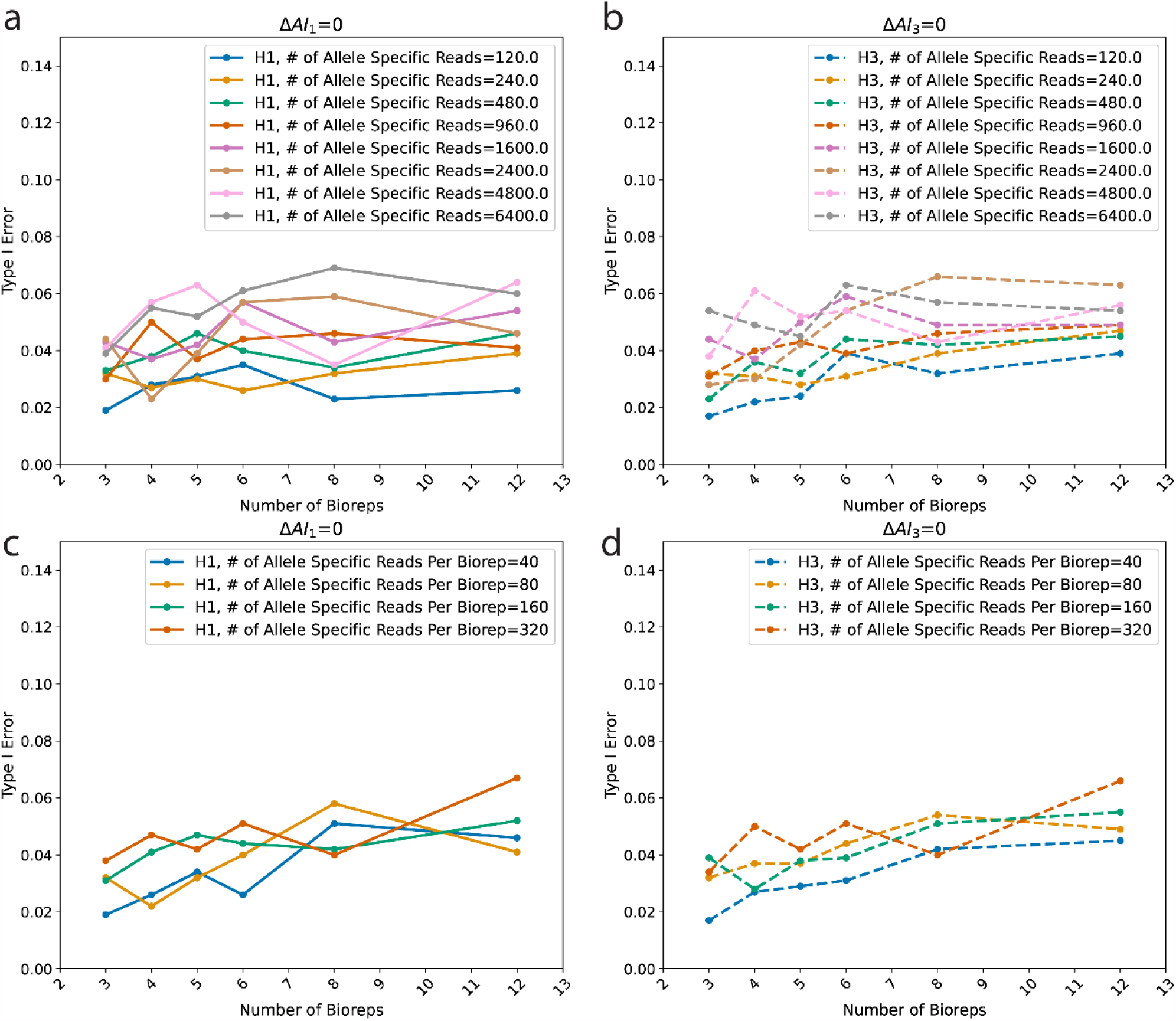
Variations in type I error (y-axis) are shown as a function of the number of biological replicates, or bioreps (x-axis) assuming different numbers of allele specific reads. H1 and H3 refer to the null hypothesis of allelic balance within a condition (H1) and the null hypothesis of equal levels of AI between the two conditions (H3). The Type I Error (y-axis) is computed as the proportion of simulations for which the Bayesian evidence against allelic balance within a condition or equal AI between conditions is < 0.05. Plots a and b show eight simulated values of the number (#) of allele specific reads, which is the sum of the reads that map unambiguously to an allele in the experiment. Plots c and d show four simulated values of the number (#) of allele specific reads per bioreps, which is the number of allele specific reads divided by the number of bioreps. Δ*AI* is deviation from the null, *i*.*e*. deviation from allelic balance in condition (Δ*AI*_1_) or the relative difference in the levels of allelic imbalance between the two conditions (Δ*AI*_3_)The probability of an allele specific read is *r*_*i,g1*_ = *r*_*i,g2*_ *= 0*.*8* and there are 1000 simulations.

Under all conditions Type I error is low (Figure 2, S1), and only exceeds the nominal value of 5% in scenarios with very high numbers of allele specific reads (n > 2400) and biological replicates.

Power (Figure 3) for detecting small deviations from the null (0.1) is less than 0.4 when the number of bioreps is 3 and only exceeds 0.6 when the number of bioreps is greater than 6 and the total number of allele specific reads is large. H1 is rejected with power > 80% when the total number of reads is at least 2400, and the number of independent biological replicates is at least 12 (an average of 200 allele specific reads per biological replicate). Power for rejecting H3 is, as expected, lower than H1 (Figure 3, S4). For Δ*AI =* 0.2 (central panels) and 960 informative reads, H1 is rejected with power > 80% with 3 biological replicates (average of 320 reads per replicate). H3 is rejected with power > 80% with 960 informative reads in 8 replicates (average of 120 reads per replicate). When Δ*AI =* 0.3 (bottom panels), power approaches 100% except when the number of informative reads is low (120). As expected, the number of simulations does not affect estimates of power (Figure S2). Power for the test of H3 for 3 biological replicates is maximal at 640 informative reads (Figure S3). When Δ*AI =* 0.3, most scenarios have power greater than 80% for both H1 and H3 (Figure 3, Figure S4). When Δ*AI* is 0.5, power for both H1 and H3 is ∼100% even when the total number of informative reads is low. This represents an extreme scenario, but one that is often observed in situations with loss of heterozygosity, indicating that in these scenarios relatively few reads are needed to detect AI with confidence (Figure S5).

**Figure 3.**
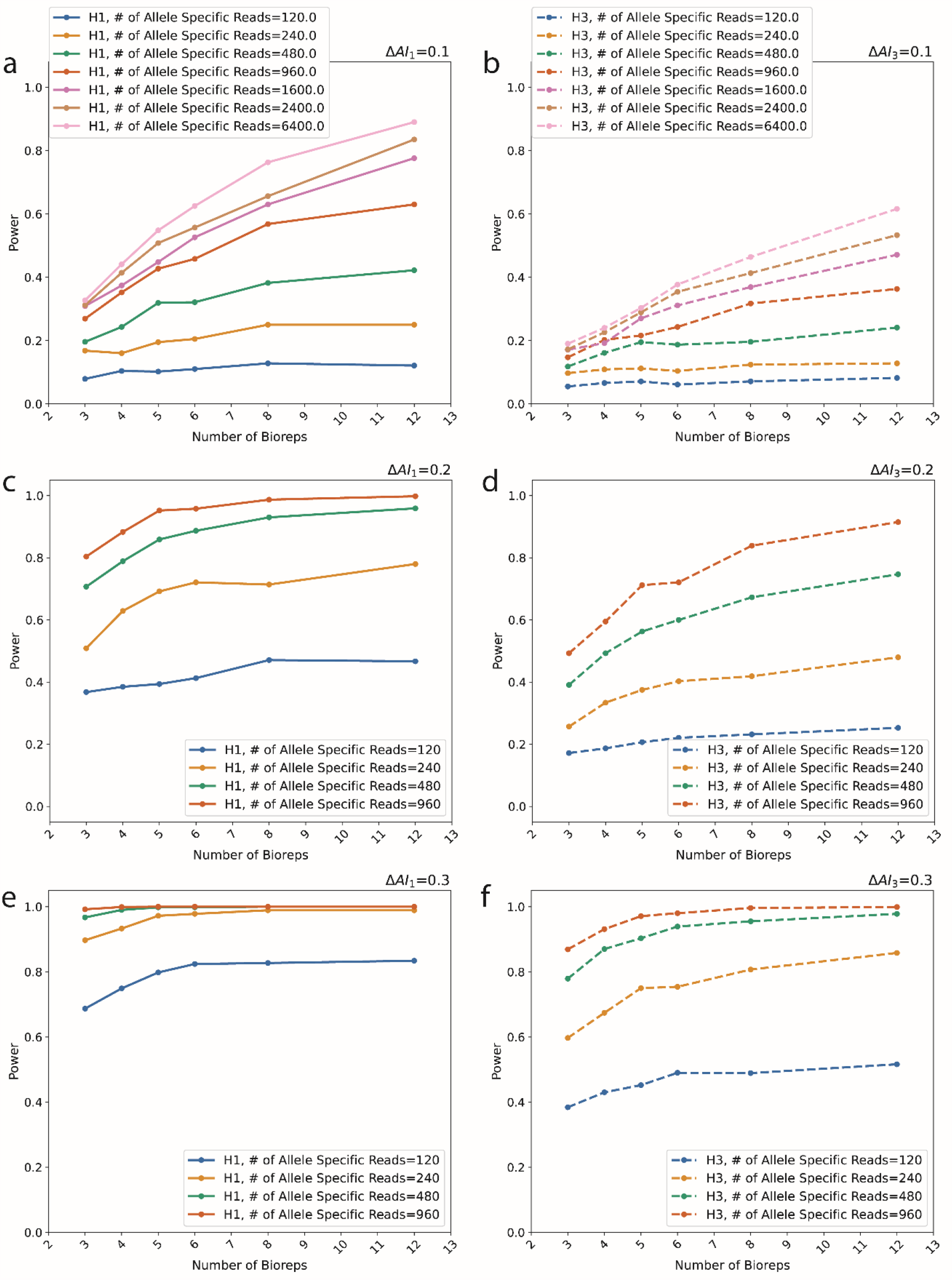
H1 refers to simulations under the alternative hypothesis of allelic imbalance within a condition and H3 refers to unequal levels of AI between the two conditions. For H1, the x-axis is the effect size, which is the relative deviation from allelic balance 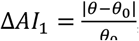, where *θ*_0_ *=* 0.5. For H3, the x-axis is the relative difference in levels of AI between the two conditions 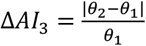 where the first condition is simulated under the null hypothesis and the second under the alternative hypothesis *θ* ≠ 0.5. The power (y-axis) is computed as the proportion of simulations for which the Bayesian evidence against allelic balance within a condition or against equal levels of AI between conditions is < 0.05. There are 1000 simulations and the probability of an allele specific read is *r*_*i,g1*_ = *r*_*i,g2*_ *= 0*.*8*. Simulations for 3, 4, 5, 6, 8, and 12 biological replicates (bioreps, x-axis) for varying numbers (#) of allele specific reads are reported.

## Discussion

Type I error rarely exceeds the nominal value of 5% even for very high numbers of allele specific reads, while increasing the number of allele specific reads substantially increases power. These observations are in agreement with other approaches [5, 15, 34, 35], including with simulations performed using a previous version of this model [19]. However, except for Zou *et al*. [5], the power of these approaches was not assessed for jointly changing the number of allele specific reads and biological replicates. BayesASE can directly test for a difference in AI between two conditions or genotypes and, accordingly, we can assess how variation in both the number of replicates and reads affects the power to not only detect AI in a condition but differences in AI between conditions.

BayesASE has adequate power to detect moderate deviations from the null hypothesis. However, the minimum number of reads and biological replicates to achieve this power is greater for smaller deviations from the null. Our simulations suggest that a minimum of 2400 informative reads across 12 replicates, 480 informative reads across 5 replicates (or a minimum of 3 replicates with a total of 960 informative reads), and 240 informative reads across 3 replicates results in >80% power to detect Δ*AI*_1_ (or Δ*AI*_2_) = 0.1, 0.2, and 0.3, respectively. While the power to detect Δ*AI*_3_ = 0.1 does not surpass 60% in our simulations, we can detect a difference in AI between conditions (Δ*AI*_3_) of 0.2 and 0.3 with comparable power for the same deviation from the null within a condition with the same number of informative reads but only when spread over more replicates (*i*.*e*. 8). A deviation from the null of Δ*AI* = 0.3 has power > 80% in most scenarios and even higher deviations can be detected with almost 100% power. Such large differences are indicative of loss of heterozygosity as observed in cancers [17] and imprinting [9, 10].

We present results of an extensive simulation study to quantify type I error and power in detecting AI using the model implemented in the BayesASE pipeline [7]. Both number of reads and number of replicates are important, and they both should be maximized. However, for any given number of reads, the best idea is to maximize the number of replicates. This is in agreement with previous studies that suggested that increased biological replication should be favored over increased depth of coverage [5, 36]. This of course should be balanced against the fact that having several replicates is more expensive. This said, we do not recommend designing any biological experiment with less than three biological replicates.

## Limitations

This simulation study, like most such studies, makes simplifying assumptions for computational ease and efficiency. It is performed under optimal scenarios for a single gene and, thus, may not account for all limitations that are inherent to real data. Thus, the recommendations based on the simulation results should be considered a minimum threshold for study size planning. However, despite their drawbacks, simulations are necessary because it is not possible to estimate power without them.

## Availability of data and materials

This study was performed using programs written in Python and R that are available using the MIT license as the package BayesASE_power: https://github.com/McIntyre-Lab/BayesASE_power. The package requires the installation of BayesASE available on PyPI: https://pypi.org/project/BayesASE/.

## Notes

https://github.com/McIntyre-Lab/BayesASE_power

### Competing Interest Statement

The authors have declared no competing interest.

