## Supplementary Methods and Figures for "Power Calculator for Detecting Allelic Imbalance Using Hierarchical Bayesian Model"

#### Model description

Let *g1* and *g2* be the two alleles of a diploid individual, respectively. For each gene or gene region, condition *i* and biological replicate (biorep) *k*, $x_{i,k}$ and $y_{i,k}$ are the number of reads that align better (or unambiguously) to allele *g1* and *g2,* respectively, while $z_{i,k}$ is the number of reads that map equally well (or ambiguously) to both alleles (Table S1).

**Table S1.** The expected number of reads ($\mu$) aligning better to allele *g1* than *g2*, $x_{i,k}$; better to allele *g2* than *g1*, $y_{i,k}$; or ambiguously, that is equally well to both alleles, $z_{i,k}$.

| $x_{i,k}$ | $y_{i,k}$ | $z_{i,k}$ |
|---|---|---|
| $(1/\alpha_i)\beta_{i,k}r_{i,g1}$ | $\alpha_i\beta_{i,k}r_{i,g1}$ | $[(1-r_{i,g1})/\alpha_i + (1-r_{i,g2})\alpha_i]\,\beta_{i,k}$ |

One important parameter in determining AI is the ability to correctly assign reads to an allele given that the read originated from that allele. We express this as the quantity $r_{i,g1}$ ($r_{i,g2}$), which is the probability of a read aligning to allele *g1* (*g2*) given that it came from that allele. Low values of these probabilities correspond to a high degree of ambiguously mapped reads, which occurs when there is little sequence divergence between the two alleles.

AI in condition *i* is measured by the parameter $\theta_i$ representing the proportion of reads originating from the allele *g1*, which that can be written as follows:

$$\theta_i = \frac{\mathbb{E}(x_{i,k}/r_{i,g1})}{\mathbb{E}(x_{i,k}/r_{i,g1} + y_{i,k}/r_{i,g2})} = \frac{1/\alpha_i}{\alpha_i + 1/\alpha_i}$$

Notably, when $\theta_i$ is close to 0 , we have one extreme case of AI with all the reads originating from $g2$. When $\theta_i = 0.5$, we have perfect allelic balance with 50% of the reads from each allele. With $\theta_i = 1$, we are in the opposite direction of extreme AI with all the reads originating from $g1$. $\theta_i$ is a function of $\alpha_i$, which is also a measure of AI representing the ratio of reads mapping to $g1$ over the reads mapping to $g2$. Consequently, $\alpha_i$ may vary from zero, when all reads map to $g2$, to infinity, when all reads map to $g1$. In the case of allelic balance, $\alpha_i = 1$.

Finally, the model allows incorporating biological variability across conditions and replicates via the variable $\beta_{i,k}$. The ideal case is $\beta_{i,k} = 1$ for all bioreps, which indicates that each biorep has the same variance.

**Simulations**

The following null hypotheses are defined:

1. Allelic balance in condition 1, *i.e.* null $H1$: $\theta_1 = 0.5$ or equivalently $\alpha_1 = 1$.
2. Allelic balance in condition 2, *i.e.* null $H2$: $\theta_2 = 0.5$ or equivalently $\alpha_2 = 1$.
3. Level of AI is the same in both conditions, *i.e.* null $H3$: $\theta_1 = \theta_2$ or equivalently $\alpha_1 = \alpha_2$.

To test these hypotheses, three scenarios are defined (Figure 1):

1. H1, H2 and H3 are satisfied
2. H1 is satisfied, H2 and H3 are violated
3. H1 and H2 are violated, H3 is satisfied

Read counts were simulated under various scenarios assuming a negative binomial model with a dispersion of 50 and the mean (μ) defined for $x_{i,k}$, $y_{i,k}$ and $z_{i,k}$ as shown in Table S1. The full list of the simulation parameters is shown in Supplementary Table 2.

The simulations were designed varying $\theta_1$ and $\theta_2$ from 0.25 to 0.75 with step 0.05. Previous work has shown that the results for $\theta_i > 0.5$ ($\alpha_i < 1$) are symmetric to those for $\theta_i < 0.5$ ($\alpha_i > 1$) [20] so we focus here on the former set of $\theta_i$ values.

$r_{i,g1}$ and $r_{i,g2}$were simulated to vary between 0.2 and 0.8 with step 0.05.

The number of bioreps was set to 3 for most simulations. When investigating the effect of varying number of bioreps on type I and type II error, the number of replicates was varied between 3 and 12.

The total number of allele specific reads was varied from 12 to 480,000. Allele specific reads are reads that map unambiguously in the simulation. Informative reads were equally distributed across bioreps.

For simplicity, all simulations were run assuming $\beta_{i,k} = 1$ for all conditions and replicates.

**Computing type I and type II error**

Under the extended simulation scenario, type I error is defined as the proportion of simulations for which the Bayesian evidence against allelic balance is less than 0.05 when simulations were performed under the null hypothesis. Three different null hypotheses are possible, as shown in Fig. 1. Hypothesis H1 and H2 are null when $\theta_1 = 0.5$ and $\theta_2 = 0.5$, respectively. H3 is null when $\theta_1 = \theta_2$. H3 can be null even when both H1 and H2 are not null if both conditions are simulated with the same level of AI.

The power to detect AI within a condition or a difference in AI between conditions is the proportion of simulations for which the Bayesian evidence against allelic balance or equal levels of AI is less than or equal to 0.05 when simulations were performed under the not null hypothesis. Within a condition, the H1 (or H2) hypothesis is not null when $\theta_1 \neq 0.5$ (or $\theta_2 \neq 0.5$). When comparing AI between conditions, the H3 hypothesis is not null when one condition is simulated with allelic balance ($\theta_1$ = 0.5) and the other with allelic imbalance ($\theta_2 \neq 0.5$). magnitudes of deviation from the null, which are measured as $\Delta AI$. Given $\theta_0 = 0.5$ ,$\Delta AI_1 = \frac{|\theta_1 - \theta_0|}{\theta_0}$ for H1 $\Delta AI_2 = \frac{|\theta_2 - \theta_0|}{\theta_0}$ for H2, and $\Delta AI_3 = \frac{|\theta_2 - \theta_1|}{\theta_1}$ for H3. Deviations of $\Delta AI$ from the null are moderate, generally between 0.1 and 0.3, with a maximum of 0.5. The reader can easily verify using the given equations that $\Delta AI$=0.5 can be obtained, when $\theta_0$ = 0.5 and $\theta_1$ = 0.25. If $\theta_0$ = 0.5 and $\theta_1$ = 0.4, then $\Delta AI$=0.2, and so on.

### Supplementary Figures

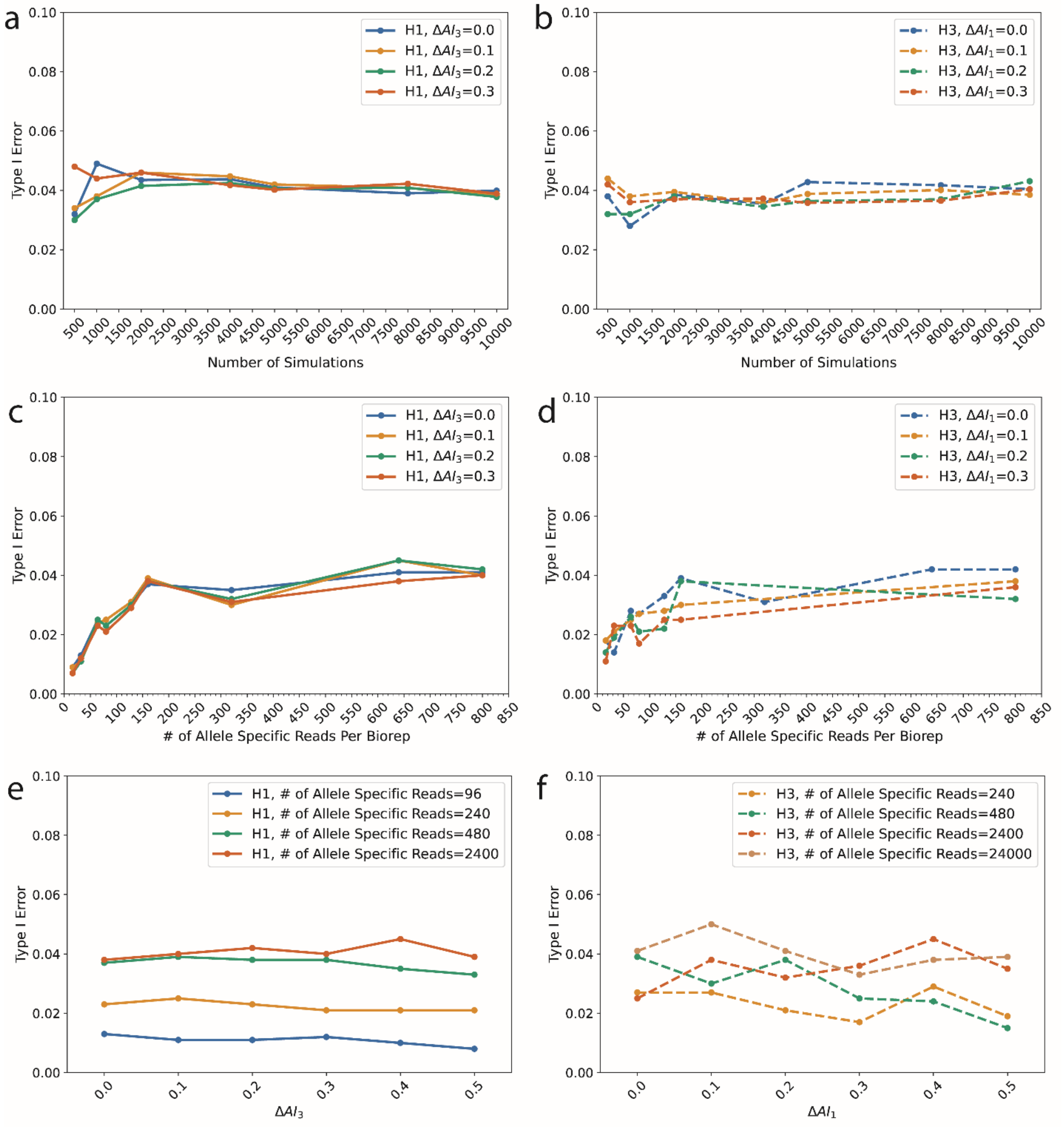


**Figure S1.** Type I error under different scenarios. (a-b) The x-axis is the number of simulations to obtain a read count dataset. The number (#) of allele specific reads was set to 2400, there were 3 bioreps, and the probability of an allele specific read was set to $r_{i,g1} = r_{i,g2} = 0.8$. Either $\Delta AI_1$ or $\Delta AI_3$ was varied from 0.5 to 0.65 by a step of 0.05 to test for H3 or H1, respectively. (c-d) The x-axis is the number (#) of allele specific reads per biological replicate (biorep). There were 1000 simulations and the probability of an allele specific read was set to $r_{i,g1} = r_{i,g2} = 0.8$. H1 and H3 were evaluated as for the simulations in a-b. (Bottom Left) H3 was evaluated using simulations of each of two conditions under the same $\theta \neq 0.5$. For H3, the effect size is the relative deviation from allelic balance in either of the two conditions $= \frac{|\theta - \theta_0|}{\theta_0}$, where $\theta_0 = 0.5$. In evaluating H1, the relative difference in the levels of allelic imbalance $\Delta AI$ was computed where the second condition was simulated under the not null hypothesis. (e-f) The x axis is the deviation from the null hypothesis of allelic balance in a condition $\Delta AI_1$ or of equal levels of AI between conditions $\Delta AI_3$. $\Delta AI_3$ was varied to test for H1 while $\Delta AI_1$ was varied to test for H3. There were 1000 simulations and the probability of an allele specific read was set to $r_{i,g1} = r_{i,g2} = 0.8$.

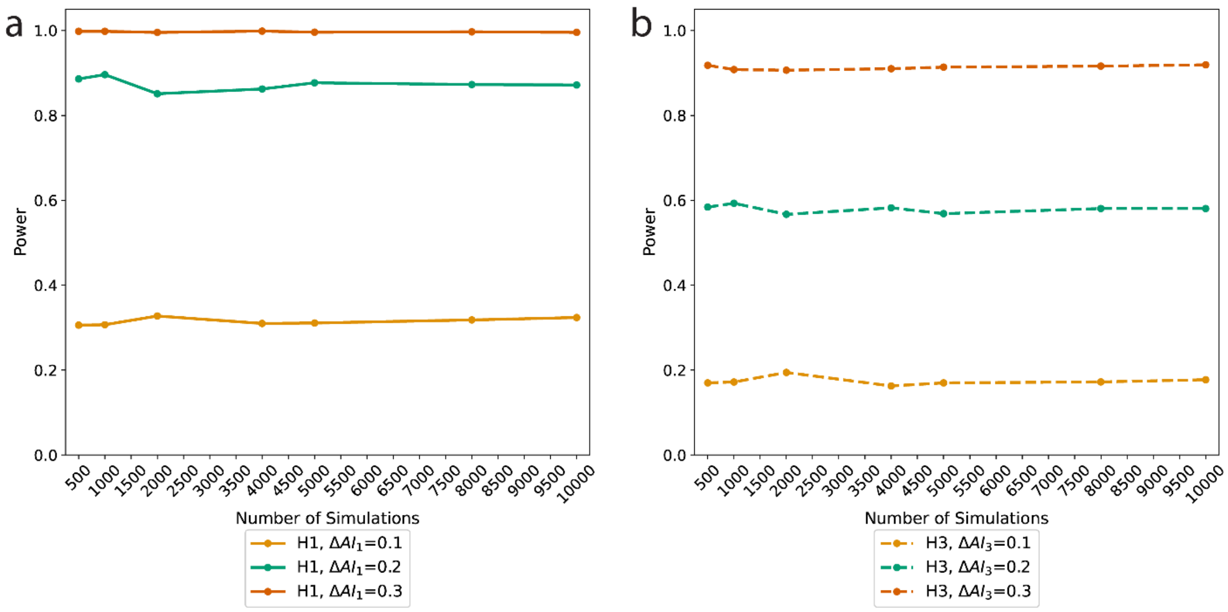


**Figure S2.** H1 and H3 refer to simulations under the not null hypothesis of allelic imbalance within a condition and unequal levels of AI between the two conditions, respectively. The x-axis is the number of simulations that were done to obtain a read count dataset. The power (y-axis) is computed as the proportion of simulations for which the Bayesian evidence against allelic balance within a condition or against equal levels of AI between conditions is $< 0.05$. In evaluating H1, the effect size is the relative deviation from allelic balance in a condition $= \frac{|\theta - \theta_0|}{\theta_0}$, where $\theta_0 = 0.5$. For H3, the relative difference in the levels of allelic imbalance $\Delta AI = \frac{|\theta_2 - \theta_1|}{\theta_1}$ was computed where the first condition and second condition were simulated under the null hypothesis of allelic balance and the not null hypothesis, respectively. The number (#) of allele specific reads was set to 2400 and the probability of an allele specific read was set to $r_{i,g1} = r_{i,g2} = 0.8$. At any given effect size or $\Delta AI$, the power to detect AI in a condition or differing levels of AI between conditions is consistent across the number of simulations.

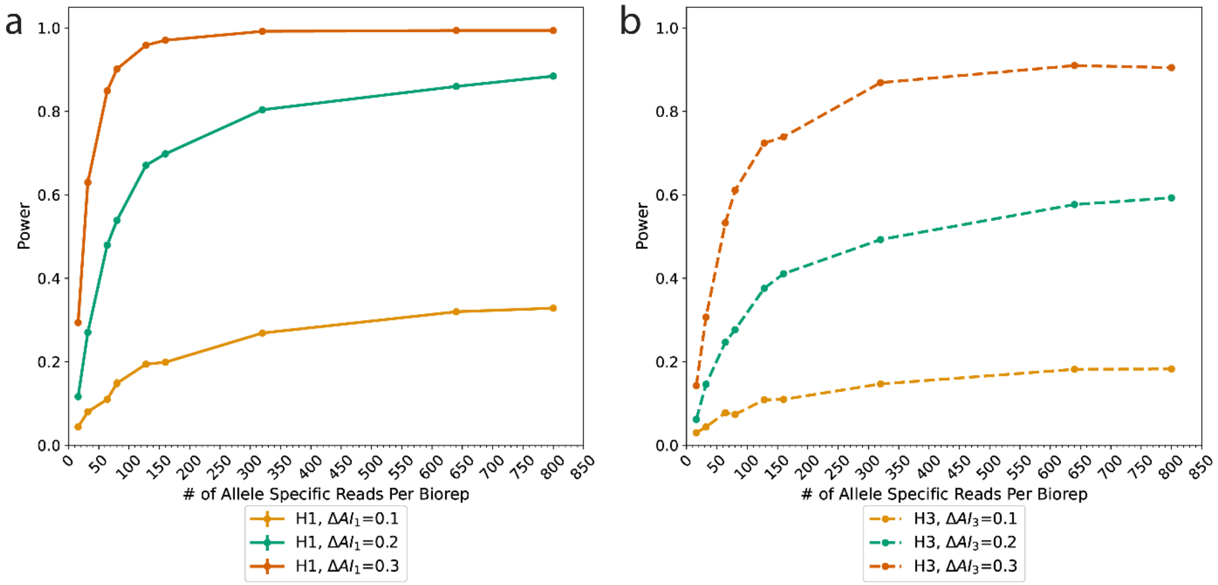


**Figure S3.** H1 and H3 refer to simulations under the not null hypothesis of allelic imbalance within a condition and unequal levels of AI between the two conditions, respectively. The x-axis is the number (#) of allele specific reads per biological replicate (biorep). The power (y-axis) is computed as the proportion of simulations for which the Bayesian evidence against allelic balance within a condition or against equal levels of AI between conditions is $< 0.05$. In evaluating H1, the effect size is the relative deviation from allelic balance in the condition $= \frac{|\theta - \theta_0|}{\theta_0}$, where $\theta_0 = 0.5$. For H3, the relative difference in the levels of allelic imbalance $= \frac{|\theta_2 - \theta_1|}{\theta_1}$ was computed where the first condition was simulated under the null hypothesis and second condition under the not null hypothesis. There were 1000 simulations, 3 biological replicates (bioreps) and the probability of an allele specific read was set to $r_{i,g1} = r_{i,g2} = 0.8$. The power to detect AI in a condition or differing levels of AI between conditions increases as the number of allele specific reads per biorep increases but does plateau for higher effect sizes and $\Delta AI$.

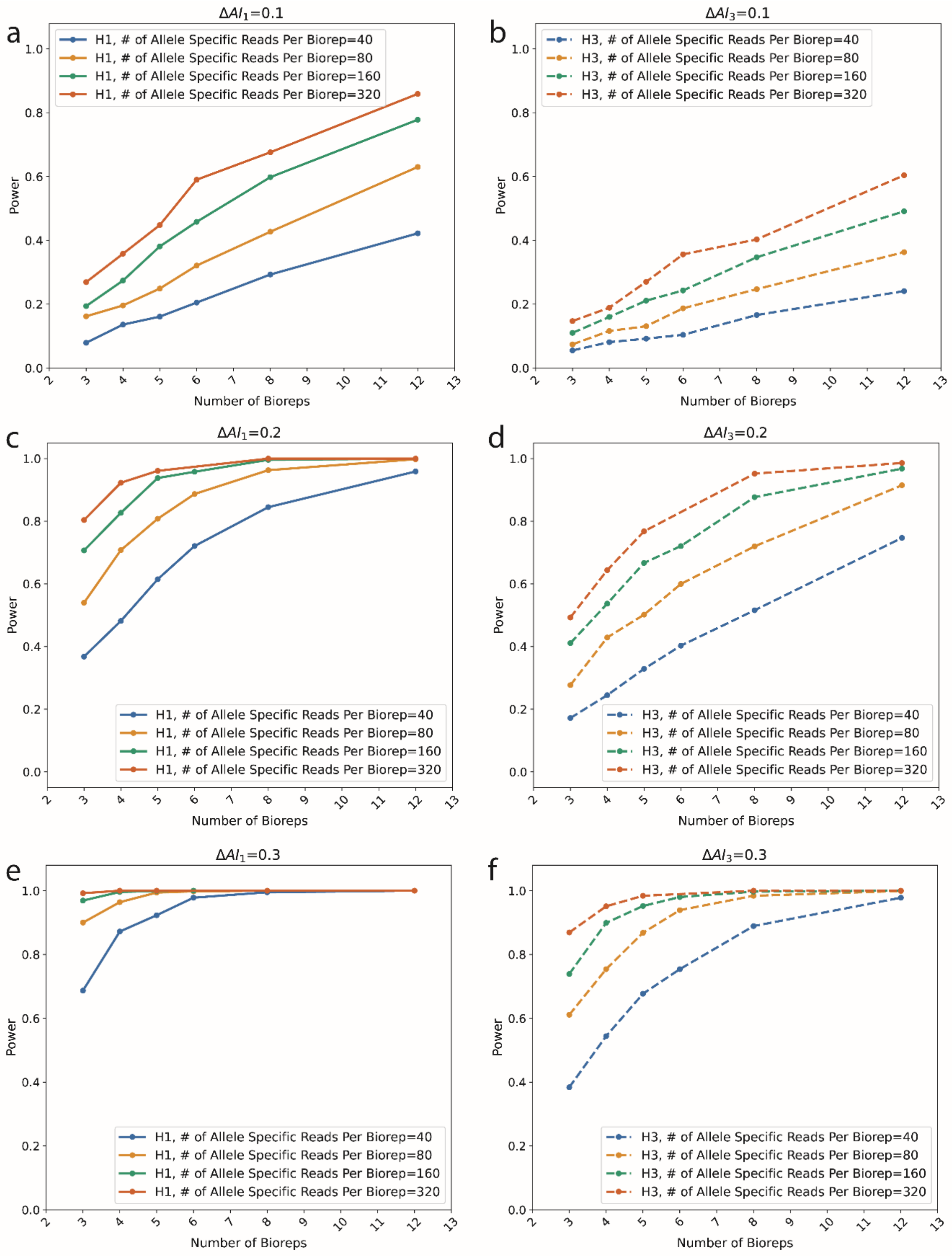


**Figure S4.** H1 and H3 refer to simulations under the not null hypothesis of allelic imbalance within a condition and unequal levels of AI between the two conditions, respectively. For evaluating H1, the x-axis is the effect size, which is the relative deviation from allelic balance in a condition $= \frac{|\theta-\theta_0|}{\theta_0}$, where $\theta_0 = 0.5$. For evaluating H3, the x-axis is the relative difference in levels of AI between two conditions $\Delta AI = \frac{|\theta_2-\theta_1|}{\theta_1}$ where the first condition simulated under the null hypothesis and the second under the not null hypothesis $\theta \neq 0.5$. The power (y-axis) is computed as the proportion of simulations for which the Bayesian evidence against allelic balance within a condition or against equal levels of AI between conditions is $< 0.05$. There were 1000 simulations and the probability of an allele specific read was set to $r_{i,g1} = r_{i,g2} = 0.8$. Simulations were performed for 3, 4, 5, 6, 8, and 12 biological replicates (bioreps, x-axis) for various number (#) of allele specific reads per biological replicate (bioreps). Increasing the number of biological replicates increases the power to detect AI or a difference in AI between conditions.

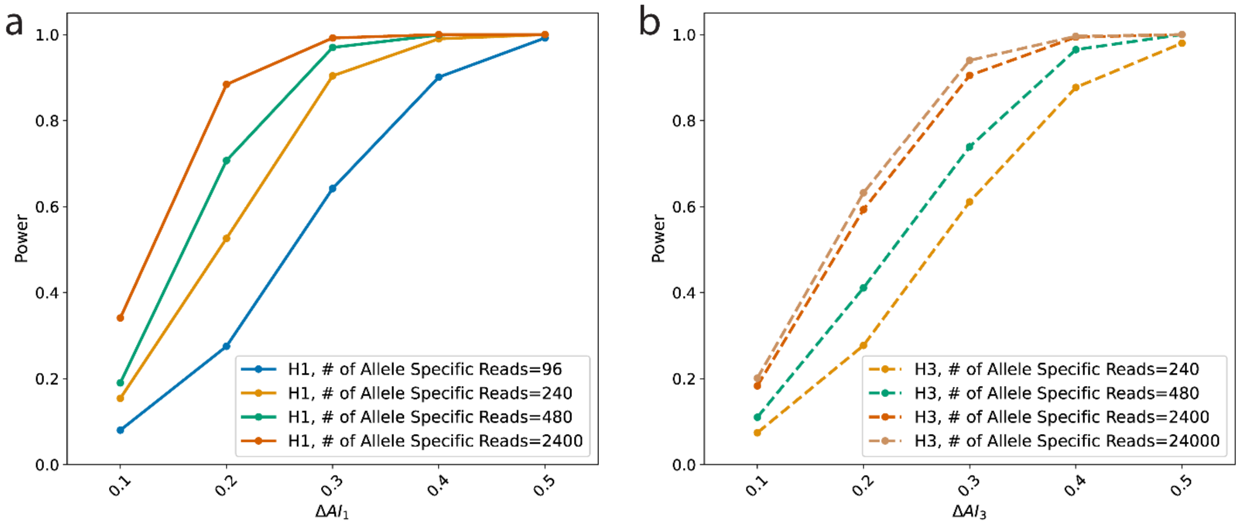


**Figure S5.** H1 and H3 refer to simulations under the not null hypothesis of allelic imbalance within a condition and unequal levels of AI between the two conditions, respectively. For evaluating H1, the x-axis is the effect size, which is the relative deviation from allelic balance in either of the two conditions $= \frac{|\theta - \theta_0|}{\theta_0}$, where $\theta_0 = 0.5$. For evaluating H3, the x-axis is the relative difference in levels of AI between two conditions $\Delta AI = \frac{|\theta_2 - \theta_1|}{\theta_1}$ where the first condition simulated under the null hypothesis and the second under the not null hypothesis $\theta \neq 0.5$. The power (y-axis) is computed as the proportion of simulations for which the Bayesian evidence against allelic balance within a condition or against equal levels of AI between conditions is $< 0.05$. There were 1000 features and the probability of an allele specific read was set to $r_{i,g1} = r_{i,g2} = 0.8$. The power to detect AI in a condition or differing levels of AI between conditions increases as the effect sizes or $\Delta AI$ increases for higher effect sizes and $\Delta AI$, but does plateau.
